# Investigating the mechanism of resistance to indaziflam in *Poa annua*

**DOI:** 10.64898/2026.07.13.738281

**Authors:** Mohit Mahey, Michael Ozolins, Joshua Miranda Teo, Marcelo L. Moretti, James T. Brosnan, Peter K. Lundquist, Eric L. Patterson

**Affiliations:** Department of Plant, Soil, and Microbial Sciences, Michigan State University, East Lansing, MI 48824, USA; Department of Biochemistry & Molecular Biology, Michigan State University, East Lansing, MI 48824, USA; Plant Resilience Institute, Michigan State University, East Lansing, MI 48824, USA; Department of Plant Sciences, University of Tennessee, Knoxville, TN 37996, USA; Department of Horticulture, Michigan State University, East Lansing, MI 48824, USA; Department of Horticulture, Oregon State University, Corvallis, OR 97331, USA

**Keywords:** *Poa annua*, indaziflam, simazine, cytochrome P450, coatamer, resistace

## Abstract

Indaziflam is an effective herbicide for controlling Poa annua, a weed, in turfgrass, orchards, and rangelands. An RNA-seq analysis was performed to identify differentially expressed genes associated with indaziflam-resistance in P. annua populations, with heterologous transformation performed for validation of candidate gene. Additionally, whole transcriptome variant calling was used to determine potential target site mutations. Transcriptome data were analyzed using weighted gene co-expression network analysis, Multiple modules were identified with high correlation to ED50 values of the resistant and susceptible populations. A cytochrome P450, CYP81A91_A, was identified that was homologous to CYP81A10 that confers resistance to five different herbicidal modes of action in *Lolium rigidum*. A single nucleotide polymorphism (SNP) causing an amino acid change (Thr432Arg) in coatomer subunit γ was identified P. annua populations and several others originating from orchards. The up-regulated and positively correlated CYP81A91_A identified, when transformed to Arabidopsis thaliana, did not provide resistance to indaziflam; however, it did provide multifold resistance to simazine, another triazine class herbicide from a different MOA. A SNP in coatomer subunit γ (Thr432Arg) was identified that may provide target site resistance given that coatomer proteins are involved in vesicle trafficking and in recycling of the cellulose synthase.

**HIGHLIGHT:** This paper is a in-depth characterization of indaziflam resistance in *Poa annua* that identifies a novel mutation in *coatomer subunit γ* as the potential root cause.

## 1 INTRODUCTION

*Poa annua* is an allotetraploid (2n = 4x = 28) species in the Poaceae family that originated from a hybridization between *Poa supina* and *Poa infirma* (Mao and Huff, 2012, 2017; Benson *et al*., 2023). This species is highly adaptable to new environments, with it being the only plant species that is found on all seven continents including Antarctica (Chwedorzewska, 2008). *Poa annua* is a highly self-pollinated species with outcrossing occurring in an estimated 0 to 15% of offspring, dependent on environmental conditions and population dynamics such as flowering synchronization, drought, nutrient availability, and air circulation (Ellis, 1973). Being allotetraploid, *P. annua* has genetic material inherited from both diploid parents (Johnson, 1995) which is hypothesized to allow it to adapt to a wider range of environments compared to either *P. infirma* or *P. supina* alone (Hovin, 1957).

During a survey conducted in 2023, *P. annua* was found to be particularly problematic in turfgrass systems (e.g., golf courses, sports fields, lawns, etc.), winter cereal grains, pastures, rangelands, and hay production (Van Wychen, 2017). A notorious weed, herbicides are the primary method of *P. annua* control; however, this heavy reliance on herbicides has led to evolved resistance in at least 12 modes of action, in some cases *P. annua* populations having multiple resistance (‘Weed Science’, 2025). Documented cases of *P. annua* evolving resistance to herbicides within several Herbicide Resistance Action Committee (HRAC) groups have been reported including: Group 2, inhibition of acetolactate synthase (ALS) (Cross *et al*., 2013; McElroy *et al*., 2013; Brosnan *et al*., 2015, 2016), Group 3, inhibition of microtubule assembly (Isgrigg III *et al*., 2002; Cutulle *et al*., 2009; Singh *et al*., 2021), Group 5, inhibition of photosynthesis at PSII (serine 264) binders (Brosnan *et al*., 2020*a*; Singh *et al*., 2021), Group 9, inhibition of 5-enol pyruvyl shikimate phosphate synthase (Binkholder *et al*., 2011; Brosnan *et al*., 2012*b*), Group 14, inhibition of protoporphyrinogen oxidase (Yu *et al*., 2018) and Group 29, cellulose biosynthesis inhibitors . Multiple and cross resistant populations of *P. annua* have been reported from Texas (Singh *et al*., 2021) to southern Australia (Barua *et al*., 2020). Evolution of multiple herbicide resistance poses a serious threat with the potential for cross-resistance to chemistry not yet on the market (Ma *et al*., 2013).

Herbicide resistance can be summarized broadly into two categories based on mechanism: target-site resistance (TSR) and non-target site resistance (NTSR). Target site resistance mechanisms affect the site of action via altered amino acid sequences (causing conformational changes reducing the ability of the herbicide to inhibit the enzyme) or changes in the expression level of the target site requiring higher herbicide concentrations to achieve effective control. Therefore, TSR mechanisms include mutations, transcript overexpression, and gene copy number variations. Conversely, NTSR is characterized by any number of mechanisms that ultimately lead to a reduction of the active herbicide available at the site of action. For example, NTSR mechanisms encompass reduced herbicide uptake and translocation, increased herbicide sequestration, and herbicide metabolism (Gaines *et al*., 2020). Among NTSR mechanisms, herbicide detoxification through metabolism mediated by cytochrome P450 monooxygenases (P450s), followed by conjugation to amino acids, glutathione, or sugars is well-recognized in plants .

Cytochrome P450 monooxygenases are one of the largest superfamilies of enzymes found in plants with 244 in *Arabidopsis thaliana* and 355 in *Oryza sativa* (Nelson, 2009). Cytochrome P450s have been classified as heme-thiolate protein-dependent oxidase systems that utilize NADPH and/or NADH to catalyze oxidation reactions including hydroxylation and demethylation^28^. They are vital for plant secondary metabolism and perform a wide diversity of reactions such as hormone synthesis, sterols and fatty acid derivatization, and the production of terpenoids. The extensive catalogue of cytochrome P450 enzymes results in overlapping protein functions and a promiscuous nature that makes them a perfect vehicle of resistance evolution (Abdollahi *et al*., 2021). P450s commonly alter small organic molecules through hydroxylation or dealkylation by utilizing high energy electrons provided by NADPH. The reduced or modified organic molecule, in this case an herbicide, is typically further inactivated by conjugation to glucose for vacuole sequestration (Kreuz *et al*., 1996) or glutathione conjugation by glutathione-S-transferase (GST) .

Resistance to triazine herbicides (HRAC Group 5) such as atrazine and simazine in *P. annua* has been well-documented globally. The first reported instance of resistance occurred in France in the early 1980s, and it has since spread extensively. In the United States, Group 5 resistance is common in *P. annua* with surveys reporting numerous resistance cases (Hutto *et al*., 2004). Similarly, widespread occurrences of Group 5 resistance have been reported across Europe and Asia, including Belgium, Germany, Japan, the Netherlands, Norway, and the United Kingdom. The primary mechanism is a specific point mutation in the chloroplast psbA gene (commonly a Ser264Gly substitution), which prevents the herbicide from binding to the D1 protein in photosystem II. While this mutation is the standard cause, NTSR has also been identified in certain populations where the plant survives through enhanced herbicide metabolism (Svyantek *et al*., 2016).

Indaziflam is a potent, triazine-like, alkylazine herbicide (HRAC Group 29) that acts as a cellulose biosynthesis inhibitor (CBI) (Brabham *et al*., 2014). Indaziflam acts preemergence by disrupting the formation of cell walls during seed germination and root cell division, causing radial swelling of root tissues and ultimately stopping root growth (Ahrens, 2015). In turfgrass management, indaziflam is typically applied in late summer or early fall as soil temperatures drop to < 21°C (the typical germination window for *P. annua*) (Taylor *et al*., 2021). Indaziflam is a highly effective option for preemergence *P. annua* control in warm-season turfgrasses like bermudagrass (*Cynodon* spp.), particularly biotypes with resistance to other modes-of-action (Brosnan *et al*., 2012*a*, 2014, 2015). Efficacy resulted in use patterns that selected for indaziflam-resistant *P. annua* populations over time, especially those that often are no longer controlled by simazine (Brosnan *et al*., 2020*b*; Pritchard *et al*., 2023; Miranda *et al*., 2026).

While simazine and indaziflam reside in different HRAC groups and employ completely different modes of action, they share similar chemical properties. At their core, they both have a central triazine ring with three side chains. It is possible that due to the promiscuous nature of cytochrome P450s, there could be overlap in the enzymes that bind with and metabolize simazine and indaziflam. Research was conducted to explore possible mechanisms of indaziflam resistance in *P. annua*, especially possible cross-resistance mechanisms affecting simazine and indaziflam.

## 2 MATERIALS AND METHODS

### 2.1 Transcriptomic analysis

#### 2.1.1 Validating phenotype of indaziflam-resistant Poa annua for RNA-seq

*Poa annua* populations used for the study were collected from golf courses throughout the southeastern United States and sent to the University of Tennessee Weed Diagnostics Center (Knoxville, TN) for resistance screening (Table 1). Seeds of these *P. annua* collections screening positive for resistance were then sent to Michigan State University (East Lansing, MI) to be included in research focused on mechanisms of resistance to indaziflam. Seeds were imbibed on moistened filter paper in the dark at 22 °C until the roots were 0.5 cm long. Different days of imbibition were conducted among populations to synchronize the root growth. Seedlings were then transplanted onto square plates containing 1.5% agar (Neogen Food Safety, Lansing, MI, USA) mixed with indaziflam at concentrations of 200, 400, 800, 1600, and 3200 picomolar. The plates were sealed with Parafilm® and placed vertically in a growth chamber under a photon flux density of 600 µmol m^-2^ s^-1^ for 16-hours daily. Seven days after exposure to indaziflam, root length was measured (Figure 1).

**Figure 1.**
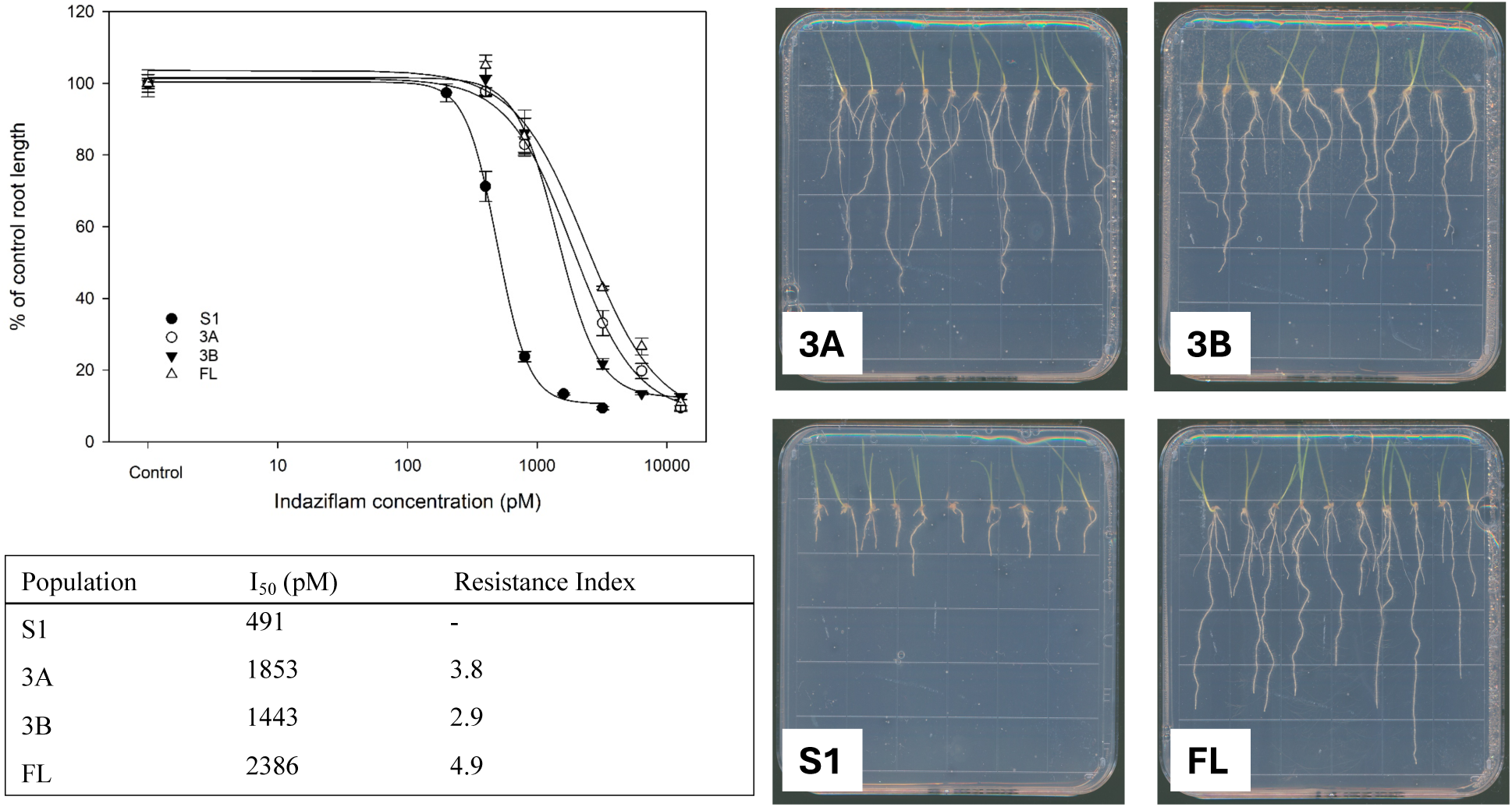
Screening of *P. annua* populations for indaziflam resistance using an agar-plate assay.

**Table 1.** Source of the *P. annua* populations, included in RNA-seq analysis, and greenhouse screening.

| **Population** | **Collection site** |
| --- | --- |
| **Site 1 (S1)** | Research plot in University Park, PA with no known herbicide resistance or history of exposure to indaziflam |
| **Site 3 (S3)** | Research plot in Knoxville, TN with no known herbicide resistance |
| **Site 5 (S5)** | A residential lawn in Wake Forest, NC that received no herbicide applications from 2005 to 2017 |
| **3A** | Golf course roughs, in Cumming, GA surviving treatment with indaziflam |
| **3B** | Golf course fairways in Cumming, GA surviving treatment with indaziflam |
| **FL** | Golf course fairways in Amelia Island, FL after surviving treatment with indaziflam |
| **MB** | Golf course fairway in Burns, TN after surviving treatment with indaziflam |
| **S18** | Golf course fairway in Thomasville, GA after surviving treatment with indaziflam |

#### 2.1.2 RNA extraction and sequencing

RNA was extracted from untreated, 4-day-old seedlings *of P. annua* using only root tips. Root tips from ten individual plants were pooled together using three susceptible (S1, S3, S5) and three resistant (FL, 3A and 3B) collections of *P. annua* (Table 1). The RNA was extracted using an RNeasy Plant extraction kit (Qiagen Inc., Hilden, Germany) and sequenced using Illumina NovaSeq6000 paired end technology (Novogene Co., Ltd., CA, USA).

#### 2.1.3 Data processing and analysis

Raw FASTQ files were filtered and trimmed for low-quality reads, and sequencing adapters were removed using *fastp* (v0.23.2) (Chen *et al*., 2018) with default parameters, including the “--trim_poly_g” flag to eliminate poly-G tail artifacts. Filtered reads were aligned to the *P. annua* reference genome (Robbins *et al*., 2022) using *HISAT2* (v2.1.0) (Kim *et al*., 2019) with default alignment settings. All analyses were performed using high-performance computing cluster (HPCC) resources provided by Michigan State University. Post-alignment, *SAMtools* (v1.19.2) (Danecek *et al*., 2021) was used to convert SAM to BAM format, followed by indexing of BAM files for downstream analyses. Read quantification was performed using *featureCounts* (v2.0.6) (Liao *et al*., 2014) to generate raw gene-level count matrices based on paired-end read data. Summary reports were compiled using *MultiQC* (v1.14) (Ewels *et al*., 2016).

#### 2.1.4 Differential gene expression

Differential gene expression was carried out via *EdgeR* (v4.2.2) (Robinson *et al*., 2010) using quasi-likelihood functionality and a generalized linear model. Comparisons were made between susceptible (S1, S3 and S5) and indaziflam-resistant populations (3A, 3B and FL) using contrast analysis. Principal component analysis and hierarchical sample clustering were utilized for identifying within sample variance (Figure 2). The p-values were adjusted by performing a multiple testing correction using the Benjamini-Hochberg method . An absolute Log2Fold Change value of > 2 and FDR-adjusted p-value of < 0.01 were used as criteria for determining up- and down-regulated genes. For visualizing sample variance, a principal component analysis (PCA) was carried out using *DESeq2* (v1.44.0) (Love *et al*., 2014). The counts were variance stabilized (Anders and Huber, 2010) and normalized using the median-of-ratios method. The normalized counts were further filtered by retaining counts of ≥ 10 across more than 90% of samples. Final PCA was plotted using ggplot2 (Wickham, 2016) package in R (v4.4.0) (R Core Team, 2024).

**Figure 2.**
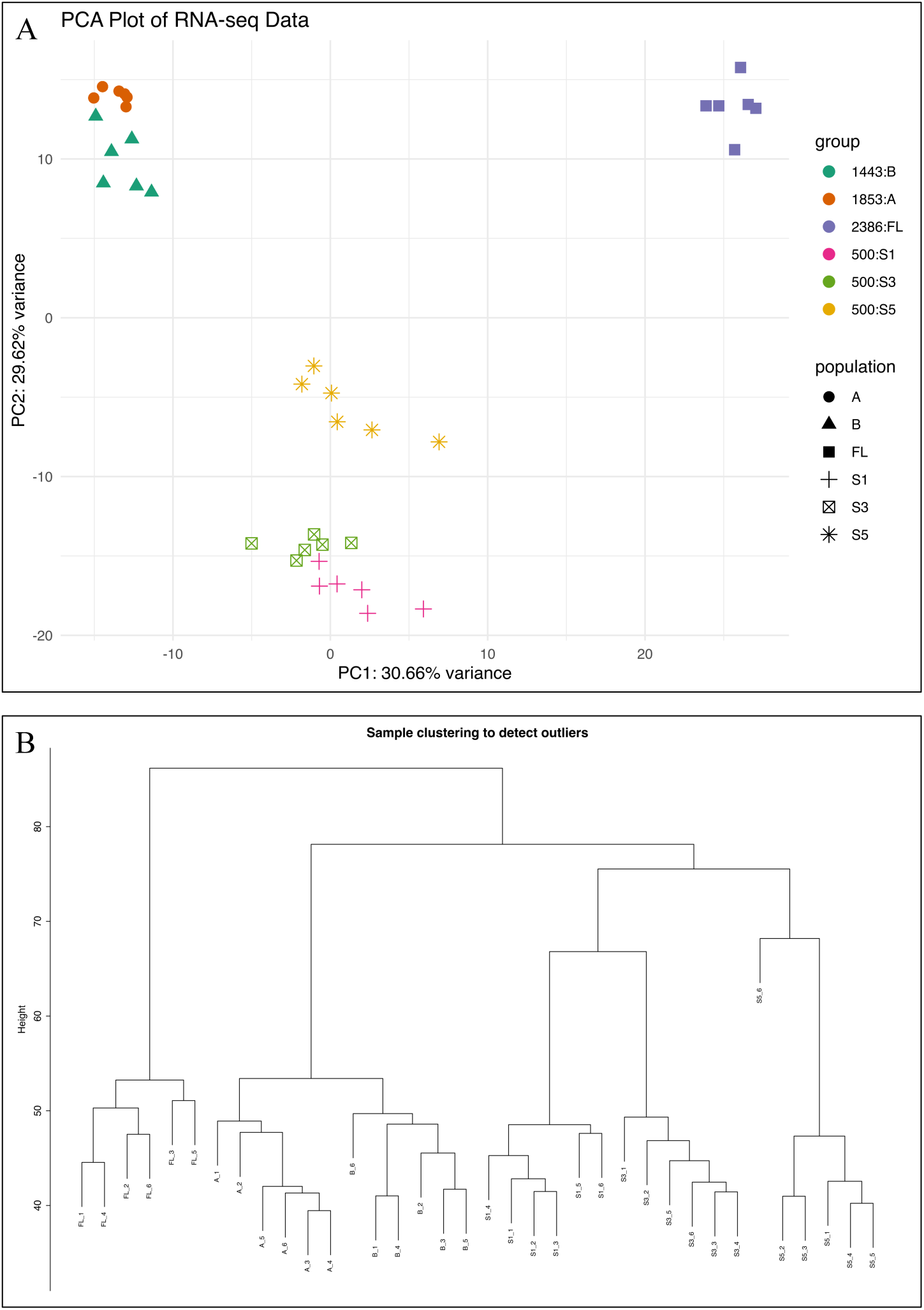
(A) Principal component analysis of the transcriptome of roots for indaziflam-resistant and - susceptible *P. annua.* The resistant and susceptible populations comprise PC1 (representing 30.66% variance), whereas PC2 is comprised of an indaziflam-resistant population from FL, as well as resistant populations 3A and 3B (representing 29.62% variance). (B) Hierarchical clustering of the samples to identify outliers. All the replicates cluster together for various populations, with no outliers.

#### 2.1.5 Weighted gene co-expression network analysis (WGCNA)

Gene expression was normalized using the *DESeq2* (v1.42.1) (Love *et al*., 2014) package. Normalized counts < 10 in 90% of samples were also filtered to remove any biologically insignificant genes. In total, 35,282 genes were selected for downstream analysis after normalization and filtering. Weighted gene co-expression network analysis (WGCNA) (Langfelder and Horvath, 2008) was conducted in R (v4.3.2) . In this analysis, soft threshold power was estimated using a scale-free topology fit index and soft power of 12 was selected. An adjacency matrix was made using normalized gene counts using a soft power of 12 and type “signed”. Further, topological overlap matrix (TOM) similarity was calculated with signed topological overlap matrix type. Minimum module size was set to 30 and mergeCutHeight to 0.20. Modules significantly associated with herbicide resistance traits were identified using a Pearson correlation between eigengene expression profiles, ED_50_ values, and the binary trait of resistant or susceptible.

In modules exhibiting significant correlation with physiological traits, hub genes were defined based on two metrics: gene significance (GS), calculated as the Pearson correlation between individual gene expression levels and physiological traits; and module membership (MM), determined by the Pearson correlation between each gene’s expression profile and the corresponding module eigengene. Genes with both GS and MM values > 0.8 were designated as hub genes (Bizouerne *et al*., 2021). After module generation, and merging modules based on a cut height of 0.20, we identified 53 distinct modules (Figure 3A).

**Figure 3.**
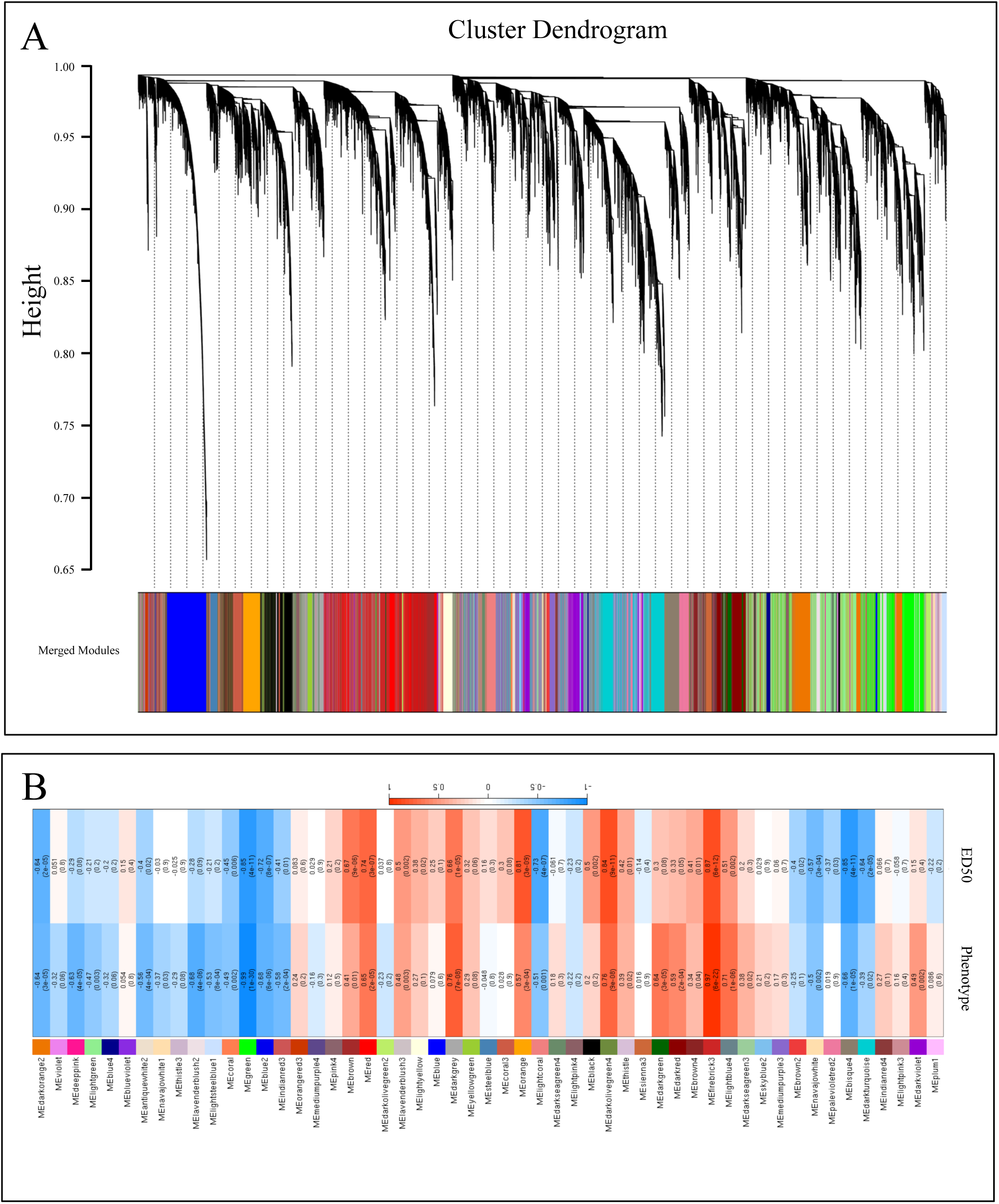
Weighted gene co-expression analysis (WGCNA) of transcriptome of *P. annua* root tips of indaziflam-resistant and -susceptible populations. (A) Cluster dendrogram showing hierarchical relationship. (B) Identification of modules associated with the resistance phenotype and ED_50_ values to indaziflam. The upper and lower values represent Pearson correlation coefficient (r) and *p-value*. The scale corresponds to the Pearson correlation coefficient (r).

### 2.2 Exploring potential for cytochrome P450 metabolism as a resistance mechanism

#### 2.2.1 Greenhouse study

Greenhouse experiments were conducted at Michigan State University (East Lansing, MI) from July 2025 to August 2025 to determine if malathion application could change the resistance spectrum of the *P. annua* populations screening positive for resistance via an agar bioassay used at the University of Tennessee Weed Diagnostics Center (Pritchard *et al*., 2023). A total of eight *P. annua* populations were included in these experiments – five resistant lines (FL, 3A, 3B, S18, and MB), as well as three susceptible lines (S1, S3, S5). Seeds were germinated in square flats filled with greenhouse media (SUREMIX, Michigan Grower Products, INC™, Galesburg, MI) and transplanted to pots of 10 cm x 10 cm x 12.7 cm. Plants were grown at a temperature range of 25 (±3)°C and a 12-hour photoperiod.

A full dose response assay was conducted using 0.25x, 0.5x, 1x (label), 2x, 4x, and 8x doses of herbicides from several mode of action groups: including mesotrione (Callisto^TM^, Syngenta Crop Protection, NC, USA), foramsulfuron (Revolver^TM^, Envu, NC, USA), and simazine (Simazine, Drexel Company, TN, USA). All herbicide treatments were mixed with non-ionic surfactant (IMPROVE^TM^ Garrco products Inc, IN, USA) at 0.25%. These herbicide applications were applied alone and in conjunction with malathion (Spectracide^TM^, Spectrum Group, MO, USA) at 2000 g a.i. ha^-1^. Malathion applications were made two hours before the application of each herbicide. All treatments were applied to plants averaging 7 cm in height using a Generation 4 DeVries Research track sprayer (DeVries Manufacturing, INC™, USA) calibrated to deliver 187 L ha^-1^ at 207 kPa. Resistant and susceptible *P. annua* populations were subjected to an increasing dose of each herbicide to determine shifts in the rate required to cause a 50% reduction in biomass (ED_50_).

#### 2.2.2 Statistical analysis

Plants were harvested at 24 days after herbicide treatment, with biomass dried for two weeks in an oven at 55°C (to remove any residual water content) and weighed. Biomass reductions were calculated by subtracting biomass of treated plants from non-treated control plants using the “drc” R package (Ritz *et al*., 2015) in R (v4.4.0) (R Core Team, 2024). *P. annua* biomass data were fit to a three-parameter log-logistic function (Equation 1) to generate dose-response curves, with and without malathion treatment.

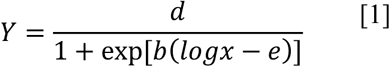

Equation 1 is the three-parameter log logistic model, where Y is the biomass reduction, x represents the herbicide dose, d is the upper limit of biomass reduction set to ≤ 100%, b is the relative slope around e, and e is the inflection point. The “*EDcomp*” function of “drc” package in R was used to compare ED_50_ values with and without malathion pre-treatment (Supplementary Table 6).

### 2.3 Identification and validating a potential candidate

#### 2.3.1 Pairwise protein alignment

WGCNA analysis identified a cytochrome P450 gene (CYP81A91_A) that was differentially expressed in all indaziflam-resistant *P. annua* populations. This gene was similar to a cytochrome P450 gene associated with multiple herbicide resistance in *Lolium rigidum* (CYP81A10v7. GenBank:QJA18355.1). To check the similarities, a pairwise protein alignment was conducted using EMBOSS needle (v6.6.0) on EMBL-EBI web service . Alignment figures were produced using Python (v3.10.15) using Matplotlib (v3.10.8) (Hunter, 2007) (Figure 4A).

**Figure 4.**
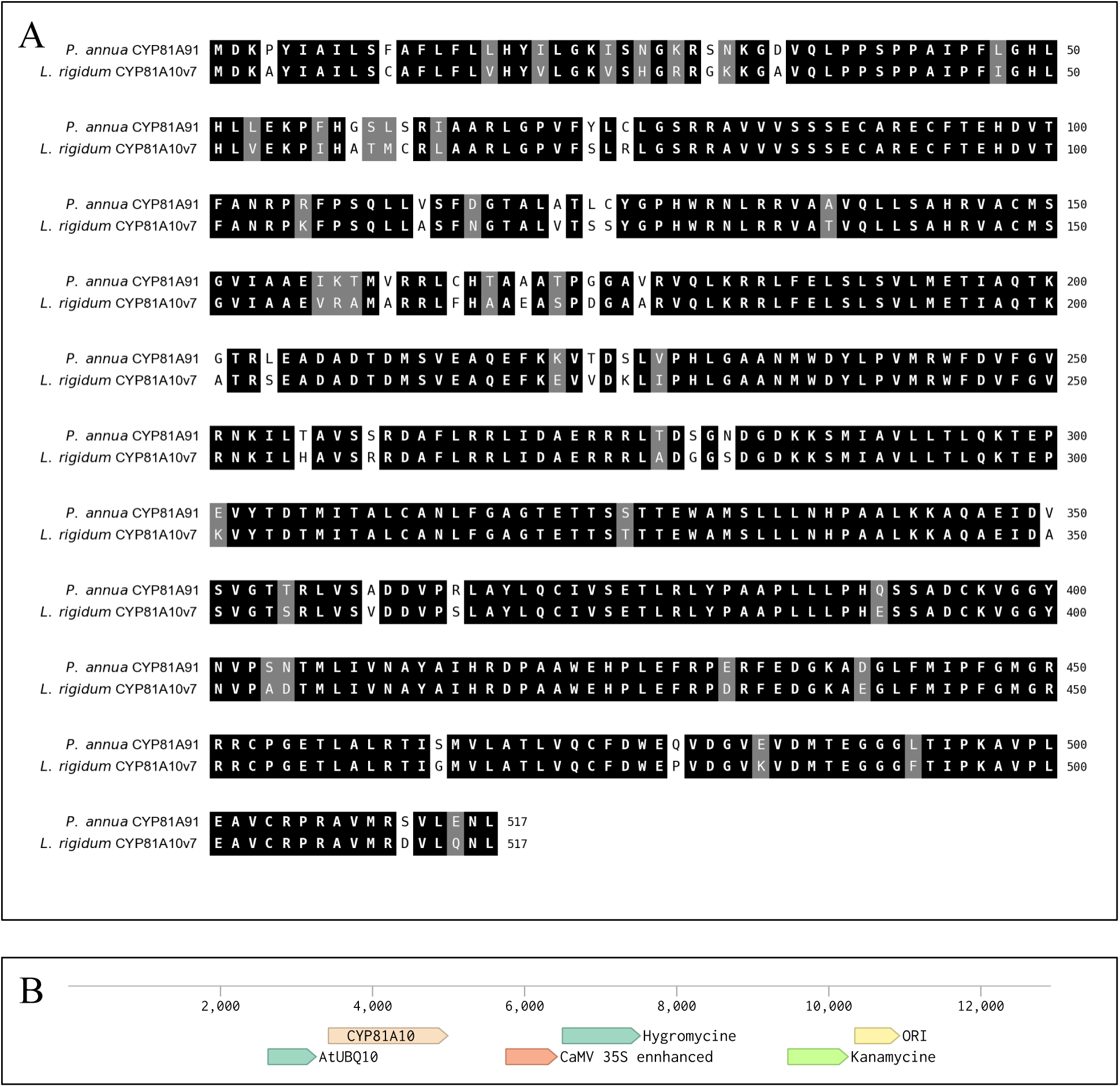
(A) Pairwise sequence alignment of *P. annua* cytochrome P450 (CYP81A91) and *Lolium rigidum* cytochrome P450 (CYP81A10v7). Alignment was performed using EMBOSS Needle . (B) Plasmid pYPQ202 transformed with *P. annua* CYP81A91_A gene with AtUBQ promoter.

#### 2.3.2 Heterologous transformation of P. annua CYP81A91_A

The potential gene candidate, CYP81A91_A, was selected for further validation using heterologous transformation. The gene was amplified from indaziflam-resistant *P. annua* populations using cDNA which was synthesized from RNA extracted from 10-day-old resistant *P. annua* seedlings. For transformation, pYPQ202 plasmid was used as a backbone (Tang *et al*., 2017). This plasmid was a gift from Yiping Qi (Addgene plasmid #86198; http://n2t.net/addgene:86198; RRID:Addgene_86198). Services from Genscript (GenScript Inc., NJ, USA) were utilized for transformation of the gene of interest.

The recombinant plasmid was first introduced into *Escherichia coli* DH5α cells via the standard heat shock transformation protocol for plasmid propagation (Thermofisher, MA, USA). The purified plasmid was subsequently transferred into *Agrobacterium tumefaciens* strain GV3101 using the freeze– thaw method (Hofgen and Willmitzer, 1988). Transformation of *Arabidopsis thaliana* was then performed using the Agrobacterium-mediated floral dip method (Clough and Bent, 1998) and redipping after one week to improve transformation efficiency. The selection of the transformants was done on plates containing 1.5% agar (Thermoscientific Inc., Waltham, MA), ½ Murashige and Skoog medium (Caisson lab science, Smithfield, UT), and hygromycin at 25 µM. Three independent transformants were selected and propagated for four generations prior to analysis.

#### 2.3.3 Verification and copy number analysis of transformation of CYP81A91_A using qPCR

Two-week-old plant leaves were collected for analysis of CYP81A91_A copy number by qPCR. The youngest leaf (∼100 mg) was collected from each plant and flash frozen with liquid nitrogen. DNA extraction was carried out using a Qiagen DNeasy Plant Mini kit (Qiagen, Hilden, Germany). The qPCR reactions were carried out in 20 µL using SYBR Green master mix (Bio-rad, CA, USA) on a bio-rad CFX96^TM^ Real-time System. The CYP81A91_A was amplified using the forward primer “ATCGACGTGTCTGTGGGTAC” and the reverse primer “GACGATGCACTGCAGGTAG”. The housekeeping gene used was GAPDH, amplified with the forward primer “TTGGTGACAACAGGTCAAGCA” and the reverse primer “AAACTTGTCGCTCAATGCAATC”. The qPCR reaction conditions consisted of an initial denaturation step of 95°C for 3 minutes, followed by 40 cycles of 95°C for 10 seconds, 60°C for 30 seconds, with a final run at 65°C to 95°C in 0.5°C increments (5 sec/step). Relative CYP abundance was estimated using comparative ΔΔCq method . All data processing and figure generation were performed using Python (v3.10.15) with NumPy (v2.2.6) (Harris *et al*., 2020), Matplotlib (v3.10.8) (Hunter, 2007), seaborn (v0.13.2) (Waskom, 2021), Claude (Anthropic, 2025) and pandas (v2.3.3).

### 2.3.4 Transformed plant screening with indaziflam

Three independently transformed *A. thaliana* (T1B6, T1C6, and T1E5) were tested for indaziflam resistance using an agar-based assay compared to a wild-type control (Col-0). 1.5% Agar plates (Thermoscientific Inc., Waltham, MA) with ½ Murashige and Skoog media (Caisson Lab Science, Smithfield, UT) were spiked with the following concentrations of indaziflam (Bayer CropScience, St. Louis, MO): 200, 150, 300, 400, and 800 picomolar (pM). The rates used were selected based on results from the initial *P. annua* screening with indaziflam (Figure 1). Seeds were surface sterilized (Lindsey *et al*., 2017) with 70% ethanol for three minutes, followed by a solution of 20% bleach (Clorox company, Oakland, CA) and 0.01% Triton X-100 (RPI Corp, Mt. Prospect, IL) for 20 minutes. Three rinsings with distilled water were used for the final cleaning. The agar plates were kept in the 4 °C cold room for 48 hours before being transferred into a growth chamber configured to provide a 25°C/18°C day/night temperature and 12 hr photoperiod delivering 150 µmol m⁻² s⁻¹ light intensity. Root length was measured seven days after indaziflam exposure.

#### 2.3.5 Transformed plant screening with simazine

Three independently transformed *A. thaliana* lines (T1B6, T1C6, and T1E5) were tested for simazine resistance compared with a wild-type control (Col-0). These lines were grown in 4 cm x 4 cm pots containing potting media (SUREMIX, Michigan Grower Products INC™, Galesburg, MI). Pots were placed inside a growth chamber for three weeks; this chamber was configured to provide a 25°C/18°C day/night air temperature and 12-hour photoperiod under 150 µmol m⁻² s⁻¹ light intensity. Simazine was applied at 0, 10, 100, 500, and 1000 g ha^-1^ using a previously described track sprayer. Plant biomass data were collected two weeks after simazine treatment, oven dried at 55°C for one week, and weighed. Biomass data were used to calculate the dose of simazine required to reduce the biomass of each *A. thaliana* line by 50% (ED_50_) and compared using “EDcomp” function of “drc” package in R (Table 5).

### 2.4 Target site mechanism discovery

#### 2.4.1 Variant calling using GATK

Whole transcriptome variant calling was done using the genome analysis toolkit (GATK) joint genotyping workflow (Brouard and Bissonnette, 2012). Filtered reads were aligned using *STAR* (v2.7.11b) (Dobin *et al*., 2013) and re-aligned again to the genome by using previously discovered splice junctions given as inputs. *SAMtools* (v1.19.2) ^42^ was used to convert SAMs to BAMs, followed by sorting and indexing the aligned reads. ReadGroups were added to the aligned reads and duplicated reads were marked using *Picard toolkit* (v3.0.0) (‘Picard’, 2025). Prior to variant calling, GATK toolkit (v4.5.0.0) (Van der Auwera and O’Connor, 2020) was used to split the reads spanning the introns and hard-clipping the overhanging sequences to make it compatible for downstream analysis. *GATK haplotype caller* was used for variant calling, using a minimum base quality score cutoff of > 20 and ignoring soft clipped bases. The variants called included single nucleotide polymorphisms (SNPs), as well as insertions and deletions (InDels). The variants were split into SNPs and InDels using values implemented in the workflow by the New York University (New York, NY USA) Genomics Core (‘NYU Genomics Core’, 2025) (read depth > 10, FS > 60, MQ < 59.9, QD < 10), with input used for base quality score recalibration (BQSR). The base recalibration was used to recalibrate the quality scores to minimize technical variation. The recalibrated reads were utilized for variant calling (using the previously described method) for downstream analysis. The variants were split into SNPs and InDels and hard filtered using consecutive single base errors (no more than 3 in 35 bp, depth more than 10, FS more than 60, MQ less than 59.9, QD less than 10). All the scripts are available within a GitHub repository (mohitmahey/whole_transcriptome_variant_calling)

#### 2.4.2 Variants filtering and annotation

The SNPs and InDels were further filtered using *bcftools* (v1.22) (Danecek *et al*., 2021) to reduce the number of biologically non-significant variants. Similar filtering was applied to both SNPs and InDels variant call files (VCFs). The compressed VCFs were normalized and variants were soft masked having genotype quality (GQ) < 20. Further, variants with minor allele frequency (MAF) of < 0.05 were removed. To remove the linked variants, pruning was done using a minimum R^2^ threshold of 0.8 within a 50 kb sliding window, to remove highly correlated variants only. The variants having a missing genotype in 90% of the populations were removed. The filtered variants were used to construct the PCA using the *adegenet* package (v2.1.11) (Jombart, 2008) in R (v4.3.2) (R Core Team, 2024) to understand the distribution of the variants in the populations. Following this, the variants specific to the indaziflam-resistant *P. annua* populations were marked and filtered using bcftools “contrast” plugin. The variants consistently present in all six replications of all indaziflam-resistant *P. annua* populations were used for further analysis. The variants were further annotated for impact using *SnpEff* (v5.2c) (Cingolani *et al*., 2012*b*) and filtered for high and moderate impact using *SnpSift* (v5.2c) (Cingolani *et al*., 2012*a*). The SNPs and InDels found were used for making biological inferences.

#### 2.4.3 Coatomer subunit γ SNP identification in Oregon populations

*P. annua* populations resistant to indaziflam were collected from orchards in Oregon^22,23^ and (Miranda *et al*., 2026)^22^ (Miranda *et al*., 2026) (Miranda *et al*., 2026)screened for the identified SNP in coatomer subunit γγ using nanopore. DNA was extracted from seeds using a ZYMO quick DNA extraction kit (ZYMO Inc., CA, USA) with RNAase treatment in between. The PCR amplification was done using Promega GoTaq green master mix (Promega, WI, USA) using a nested PCR protocol (Green and Sambrook, 2019). Two pairs of primers were used, with PCR product from the first pair used as input for the second pair. First pair primers were: forward, F1= “GCACTGCAAACGTAAACCAA” and pair one reverse, R1 = “CTTCGAAAAGCCAGAAGGTG” was used first with 95°C initial denaturing for 5 minutes, 95°C denaturing for 45 seconds, 60°C annealing for 30 seconds, extension of 72°C for 70 seconds for 34 cycles, followed by final extension of 72°C for 5 minutes. Further pair 2 forward, F2 = “ATACCTCGCAAGGCTGCAT” and pair 2 reverse, R2 = “GCCACTGGGAAGTTCTCAAA” was used for the second round following the same PCR protocol. The final product was run on a 1% agarose gel for confirmation, with DNA purification done using GeneJET Gel purification kit (ThermoScientific^TM^, MA, USA). The final DNA sequence was sent to Plasmidsaurus Inc. for sequencing and further figure generation was done using seaborn (v0.13.2) (Waskom, 2021) and Claude (Anthropic, 2025).

## 3 RESULTS

### 3.1 Agar-plate assay for pre-emergence resistance

Indaziflam-resistant *P. annua* populations originating from golf courses in the southeastern United States generated greater ED_50_ values than susceptible lines. ED_50_ values for indaziflam-resistant *P. annua* populations ranged from 1443 to 2386 pM, compared to 491 pM for the susceptible control. These responses illustrated resistance factors of 2.9 to 4.9 (Figure 1).

### 3.2 RNA quality control, filtering, and metrics

To do an untargeted assessment of gene expression differences in indaziflam-resistant and susceptible *P. annua* populations, we performed an RNA-seq experiment on 4-day-old seedlings. Raw reads ranging between 41-58 million were generated across 36 samples, after filtering for low quality reads (Supplementary Table 1). More than 99% of raw reads passed quality checks with an average of 96% reads passing the Q30 quality score. The average duplication rate was 24% across the samples (Supplementary Table 2). The alignment rate to the *P. annua* genome using HISAT2 ranged from 85% to 95%. Furthermore, *featureCounts* assigned a feature to an average of 70% of the aligned reads with average unassigned and multimap reads being ∼15%.

A PCA was performed that compared the relationship between the replicates and experimental groups to identify outliers or inconsistencies in the data. Principal component analysis revealed distinct clustering for each population, with indaziflam-resistant populations 3A and 3B clustering closely, as well as susceptible populations S1, S3, and to a lesser extent S5, clustering together as well (Figure 2A). The indaziflam-resistant population, FL, formed a separate cluster from every other population included in this experiment. The PC1 component represented 30.66% of the total variance, mostly distinguishing the indaziflam-resistant population, FL, and other two indaziflam-resistant populations (3A and 3B), while PC2 represented 29.62% of the variance and mainly separated indaziflam-resistant and susceptible populations (Figure 2A). As an orthogonal method for further visualizing transcriptome sample clustering, we utilized hierarchical clustering (Figure 2B). It revealed all indaziflam-resistant populations clustering separately from susceptible accessions. All the replicates also clustered together suggesting no outliers and consistency among replicates.

### 3.3 Weighted gene co-expression network analysis and DGE

To understand gene expression beyond pairwise comparisons and potentially identify regulatory machinery that was co-expressed within indaziflam-resistant *P. annua*, a weighted gene co-expression analysis was performed on a set of 35,282 genes expressed in the RNAseq data collected from indaziflam- resistant and -susceptible *P. annua* populations. Each module was represented by an eigengene, which is an idealized gene representing the overall expression trend (in a certain module) that was either a positively or negatively correlated to the ED_50_ value (i.e. quantifiable resistance) as well as the binary trait (resistant or susceptible) assigned to each population (Figure 3B). The modules selected for further exploration were chosen based on strong positive correlation to both ED_50_ values and binary traits (resistant or susceptible) to identify those with the most biologically relevant significance. Many modules had no or very little relation to either ED_50_ or the binary trait. We chose to focus on three modules which were very strongly correlated to both ED_50_ values and the binary trait of resistant or susceptible: MEfirebrick3 (r = 0.87), MEdarkolivergreen4 (r = 0.84), and MEgreen (r = -0.85) (Figure 3B).

The MEfirebrick3 module had the strongest positive correlation with both ED_50_ (0.87) and the binary trait (0.97), containing 1,393 transcripts, of which 75 were hub genes. The hub genes contained multiple cytochrome P450s, along with an ABC transporter (Log2Fold = 0.87) and a GST (Log2Fold = 1.11), strongly suggesting the involvement of herbicide metabolism as a mechanism of resistance in these *P. annua* populations. Among all the cytochrome P450s in the entire module (Table 2), CYP81A91_A had the highest module membership (MM) (0.93) and gene significance (GS) (0.91). CYP81A91_A was expressed 3.07-Log2fold higher in indaziflam-resistant *P. annua* compared to susceptible populations. Presence of several metabolism genes and secretory-related proteins was noted as well. Secretory carrier-associated membrane protein 4 and coatomer subunit beta-2 were present suggesting a role of the endocytosis pathway in indaziflam resistance (Supplementary Table 3).

**Table 2.**
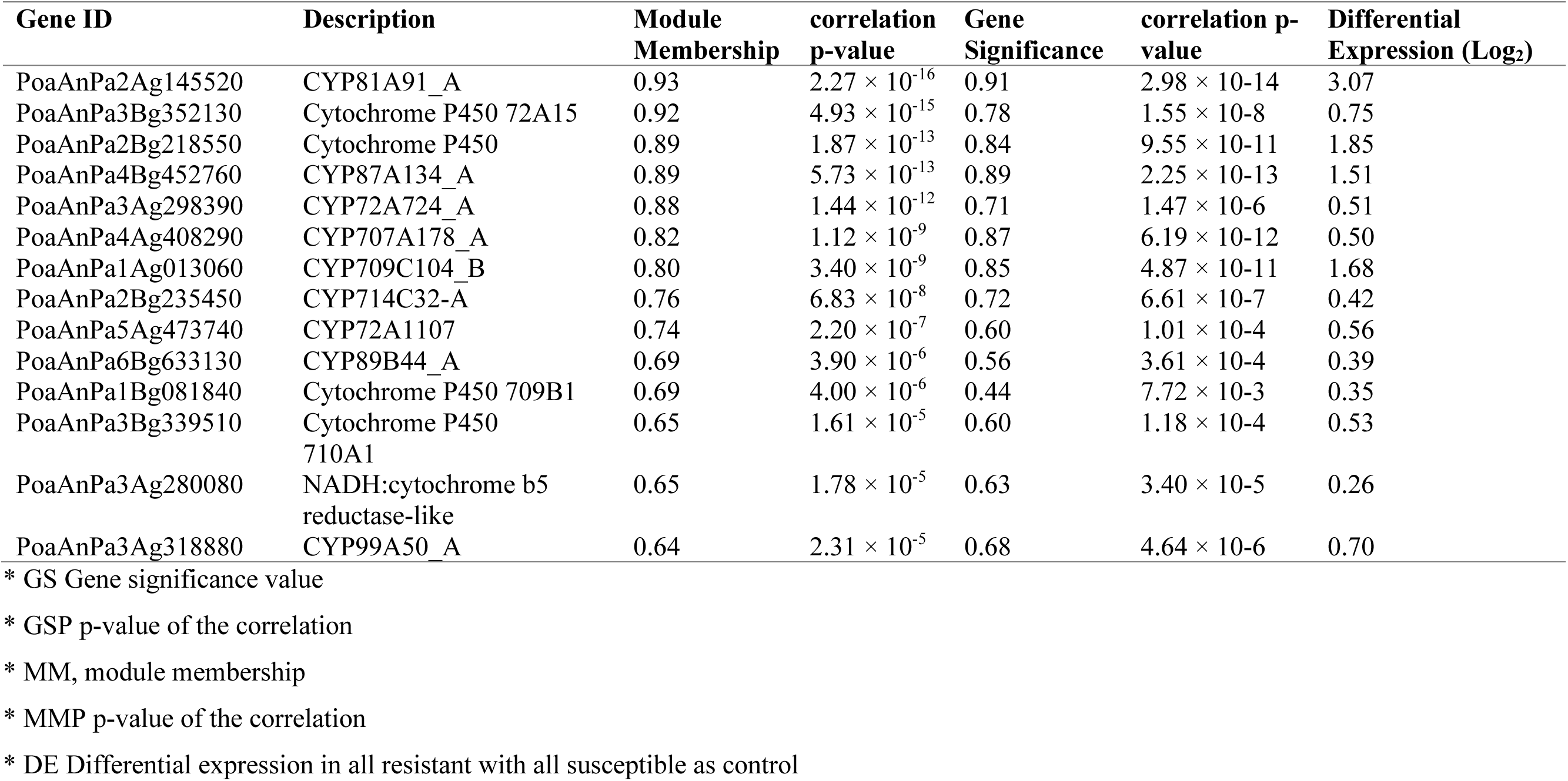
All Cytochrome P450s identified in the firebrick3 module.

| Gene ID | Description | Module Membership | correlation p-value | Gene Significance | correlation p-value | Differential Expression (Log<sub>2</sub>) |
| --- | --- | --- | --- | --- | --- | --- |
| PoaAnPa2Ag145520 | CYP81A91_A | 0.93 | $2.27 \times 10^{-16}$ | 0.91 | $2.98 \times 10^{-14}$ | 3.07 |
| PoaAnPa3Bg352130 | Cytochrome P450 72A15 | 0.92 | $4.93 \times 10^{-15}$ | 0.78 | $1.55 \times 10^{-8}$ | 0.75 |
| PoaAnPa2Bg218550 | Cytochrome P450 | 0.89 | $1.87 \times 10^{-13}$ | 0.84 | $9.55 \times 10^{-11}$ | 1.85 |
| PoaAnPa4Bg452760 | CYP87A134_A | 0.89 | $5.73 \times 10^{-13}$ | 0.89 | $2.25 \times 10^{-13}$ | 1.51 |
| PoaAnPa3Ag298390 | CYP72A724_A | 0.88 | $1.44 \times 10^{-12}$ | 0.71 | $1.47 \times 10^{-6}$ | 0.51 |
| PoaAnPa4Ag408290 | CYP707A178_A | 0.82 | $1.12 \times 10^{-9}$ | 0.87 | $6.19 \times 10^{-12}$ | 0.50 |
| PoaAnPa1Ag013060 | CYP709C104_B | 0.80 | $3.40 \times 10^{-9}$ | 0.85 | $4.87 \times 10^{-11}$ | 1.68 |
| PoaAnPa2Bg235450 | CYP714C32-A | 0.76 | $6.83 \times 10^{-8}$ | 0.72 | $6.61 \times 10^{-7}$ | 0.42 |
| PoaAnPa5Ag473740 | CYP72A1107 | 0.74 | $2.20 \times 10^{-7}$ | 0.60 | $1.01 \times 10^{-4}$ | 0.56 |
| PoaAnPa6Bg633130 | CYP89B44_A | 0.69 | $3.90 \times 10^{-6}$ | 0.56 | $3.61 \times 10^{-4}$ | 0.39 |
| PoaAnPa1Bg081840 | Cytochrome P450 709B1 | 0.69 | $4.00 \times 10^{-6}$ | 0.44 | $7.72 \times 10^{-3}$ | 0.35 |
| PoaAnPa3Bg339510 | Cytochrome P450 710A1 | 0.65 | $1.61 \times 10^{-5}$ | 0.60 | $1.18 \times 10^{-4}$ | 0.53 |
| PoaAnPa3Ag280080 | NADH:cytochrome b5 reductase-like | 0.65 | $1.78 \times 10^{-5}$ | 0.63 | $3.40 \times 10^{-5}$ | 0.26 |
| PoaAnPa3Ag318880 | CYP99A50_A | 0.64 | $2.31 \times 10^{-5}$ | 0.68 | $4.64 \times 10^{-6}$ | 0.70 |

\* GS Gene significance value
\* GSP p-value of the correlation
\* MM, module membership
\* MMP p-value of the correlation
\* DE Differential expression in all resistant with all susceptible as control

The MEdarkolivegreen4 module was also strongly positively correlated with indaziflam resistance containing 2,124 transcripts, of which 99 were hub genes. The hub genes included multiple Ca²⁺-dependent kinases, a calmodulin-like protein (CML13), scavenging enzymes (i.e. peroxidases), and R-genes (RGA2). These responses point to a module finely tuned for integrating calcium signals with transcriptional and post-translational regulation under both abiotic and biotic challenges. Metabolic enzymes (e.g. proline aminopeptidase) further suggest adjustments in osmolyte levels, also indicating that MEdarkolivegreen4 is a module associated with plant stress response (Supplementary Table 4).

The MEgreen module was negatively correlated with indaziflam resistance, containing 832 transcripts, of which 21 were hub genes. This module contained proteins mainly related to cell-wall biogenesis, protein turnover, metabolism, and transport. Relevant proteins included cell wall modification genes such as glycosyl transferase and dirigent protein. An ABC transporter B family member in this module could indicate transportation of herbicide to the vacuole. Stress-related proteins such as A1 proteases in response to oxidative stress were also found (Supplementary Table 5).

### 3.4 Identification of CYP81A91_A as potential candidate

The CYP81A91_A identified in the WGCNA analysis was chosen as a potential candidate gene for further investigation. The P450 family has been widely reported to cause herbicide resistance (Yuan *et al*., 2007), generally via phase one metabolism. The upregulation of the CYP81A91_A (along with high module membership and high gene significance in a module) provided support for CYP81A91_A as a likely candidate. Further, the protein sequence of CYP81A91_A had an overall 87.4% sequence identity (92.8% sequence similarity across a gapless region) (Supplementary Table 7) to a cytochrome P450 (CYP81A10v7. GenBank:QJA18355.1) known to confer herbicide resistance in *Lolium rigidum* (Figure 4A). This CYP81A91_A gene identified in indaziflam-resistant *P. annua* was transformed into three independent *A. thaliana* lines and its expression was driven by the AtUBQ promoter to provide consistent overexpression in all tissue types and growth stages (Figure 4B). Quantitative PCR verified the successful transformation of the *CYP81A91_A* in all three independent lines, having varied relative abundance of transformed genes (Supplementary Figure 1). The average amplification cycle of the transformed line T1B6 was 11.6, which was considerably higher than the 22.3 and 18.5 for T1E5 and T1C6, respectively (Supplementary Table 8).

### 3.5 Transformed A. thaliana response to indaziflam and simazine

We observed no significant root growth differences in three independently transformed *A. thaliana* transgenic lines compared to the wild-type on indaziflam-treated plates (Supplementary Figure 2A–F). The ED_50_ values for indaziflam in the wild-type (Col-0) and three transformed *A. thaliana* lines (T1B6, T1C6, and T1E5) were 324.32 pM, 321.75 pM, 310.65 pM, and 308.76 pM, respectively, with no statistically significant differences detected amongst lines (Table 3, Figure 5). However, responses to simazine applied postemergence were significantly different among lines (Figure 6, Supplementary Figure 3). Transformed lines (T1C6, T1B6, T1E5) had ED_50_ values of 831.33, 339.05, and 262.74 (g ha^-1^), respectively, compared to 1.01 g ha^-1^ for the wild-type control (Col-0) (Table 4).

**Figure 5.**
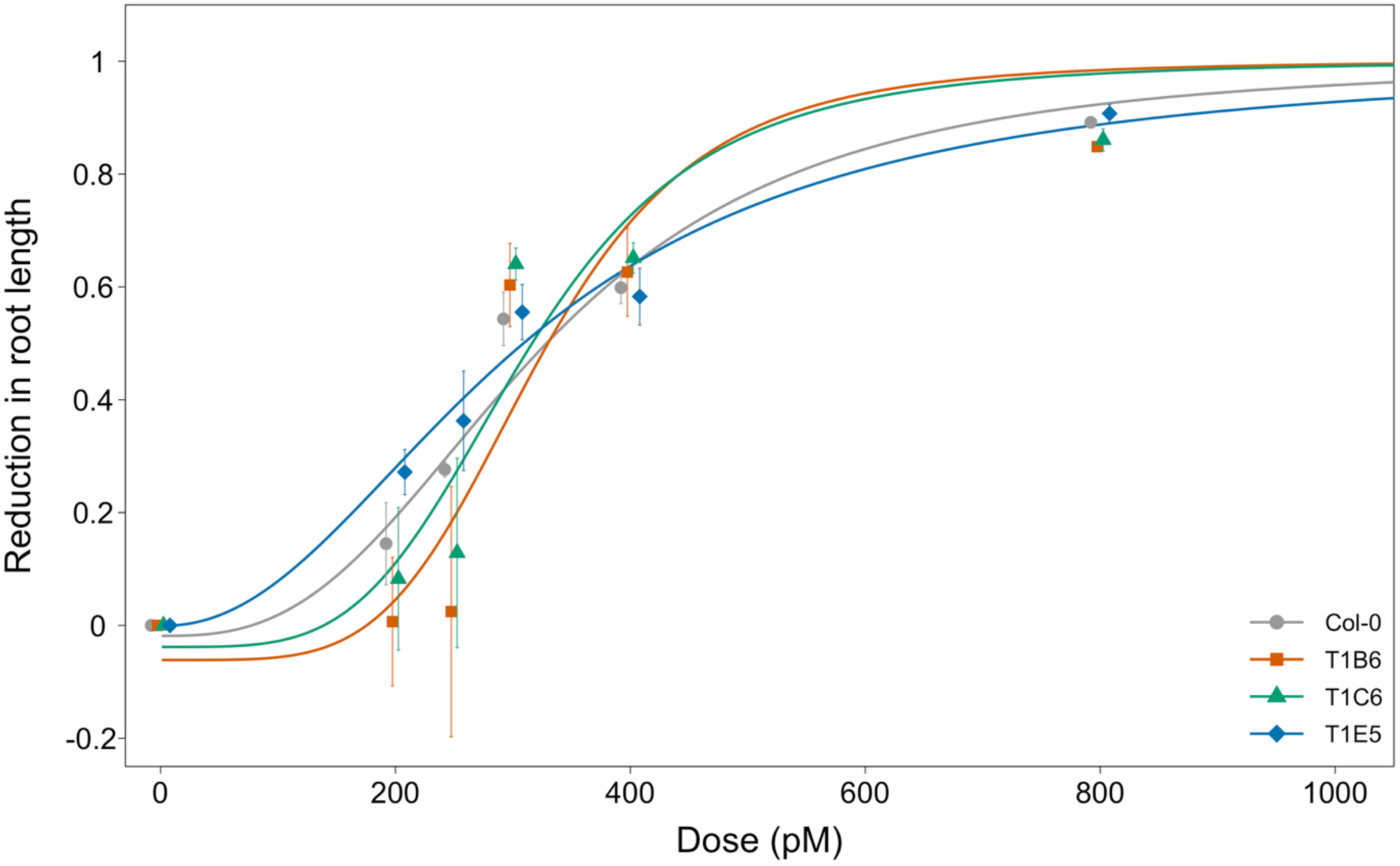
Root length of three independent transformed lines of *A. thaliana* in response to increasing doses of indaziflam. Data fit to a three-parameter logistic function (LL.3u) where the upper limit is equal to 1.

**Figure 6.**
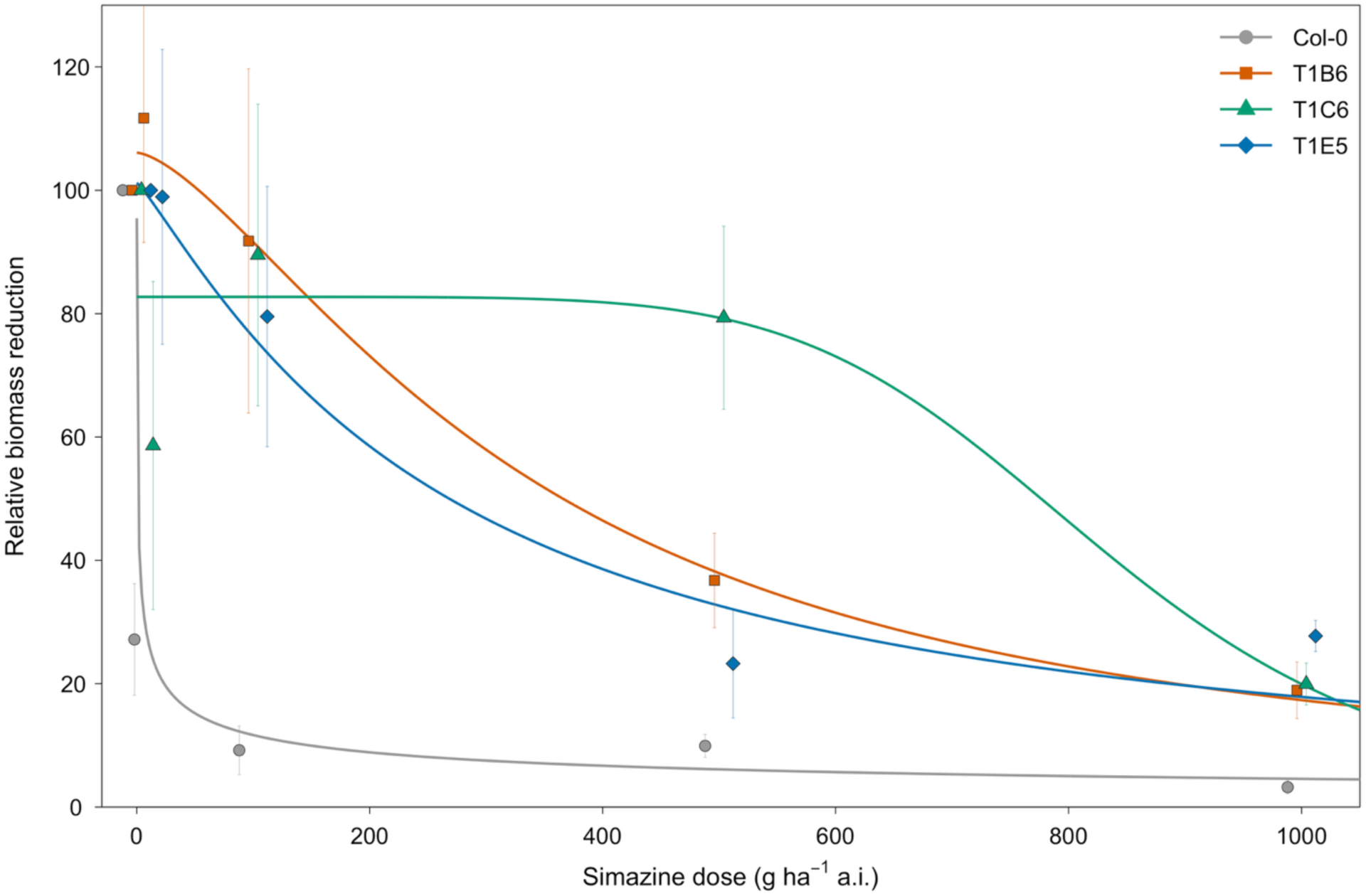
Dry biomass reductions (as a percentage of the non-treated) of three independent transformed lines of *A. thaliana* in response to increasing doses of simazine. Data fit to a three-parameter logistic function (LL.3). Error bars represent the standard error of the mean at each dose.

**Table 3.** ED_50_ values for indaziflam treated transformed *A. thaliana* plants. The table reports the estimated dose of indaziflam (pM) that reduces the root length by 50% relative to the untreated control (ED_50_), fitted using a three-parameter log-logistic model (LL.3u), where the upper limit is equal to 1.

| **Lines** | **ED<sub>50</sub> (pM)** | **Std. Error (SE)** | **95% CI Lower** | **95% CI Upper** |
| --- | --- | --- | --- | --- |
| Col0 | 324.32 | 18.00 | 285.95 | 362.70 |
| T1E5-T4 | 308.77 | 22.03 | 261.81 | 355.73 |
| T1C6-T4 | 310.66 | 27.34 | 252.39 | 368.93 |
| T1B6-T4 | 321.75 | 31.96 | 253.63 | 389.88 |

**Table 4.** ED_50_ estimates for simazine dose-response in transformed *Arabidopsis thaliana* (T1B6, T1C6, T1E5) compared to a non-transformed wild-type (Col-0). The table reports the estimated dose of simazine (g ha⁻¹) that reduces shoot dry biomass by 50% relative to the untreated control (ED_50_), fitted using a three- parameter log-logistic model (LL.3) implemented in the *drc* package.

| Transgenic Lines | ED<sub>50</sub> (g ha<sup>-1</sup>) | Std. Error (SE) | 95% CI Lower | 95% CI Upper |
| --- | --- | --- | --- | --- |
| Col-0 | 1.01 | 1.06 | -1.23 | 3.24 |
| T1B6 | 339.05 | 136.53 | 50.99 | 627.11 |
| T1C6 | 831.33 | 263.02 | 276.40 | 1386.25 |
| T1E5 | 262.74 | 120.61 | 8.29 | 517.20 |

**Table 5.** Effects of the simazine application calculated by comparison of ED_50_ values of the various transgenic lines to wild-type control, Col-0.

| Transgenic Lines | ED50 comparison | Resistance Ratio | 95% CI Lower | 95% CI Upper | p-value |
| --- | --- | --- | --- | --- | --- |
| Col-0/T1B:50/50 | 0.003 | 335.737 | -0.017 | 0.023 | $1.1 \times 10^{-75}$ |
| Col-0/T1C:50/50 | 0.001 | 822.845 | -0.007 | 0.009 | $3.0 \times 10^{-102}$ |
| Col-0/T1E:50/50 | 0.004 | 260.401 | -0.022 | 0.030 | $3.7 \times 10^{-68}$ |
| T1B/T1C:50/50 | 0.408 | 2.451 | 0.044 | 0.772 | $1.8 \times 10^{-03}$ |
| T1B/T1E:50/50 | 1.289 | 0.776 | -0.206 | 2.785 | $7.0 \times 10^{-01}$ |
| T1C/T1E:50/50 | 3.160 | 0.316 | -0.059 | 6.379 | $1.9 \times 10^{-01}$ |

### 3.6 SNP analysis in resistant individuals

From the whole transcriptome variants analysis, we detected 70,971 high-quality InDels and 289,694 SNPs across all 36 samples. We used these high-quality SNPs (Figure 7A) and InDels (Figure 7B) for a PCA to distinguish population structure. In both PCA analyses of SNPs and InDels, the indaziflam- resistant populations (3A, 3B, FL) clustered together and away from susceptible populations suggesting similar genetic structure. Among susceptible *P. annua*, each of the three populations (S1, S3, S5) separated into unique, non-overlapping clusters spread across PC2. In SNPs based on PCA (Figure 7A), the PC1 differentiated the indaziflam-resistant and susceptible populations (accounting for 21.45% variance), while PC2 accounted for 13.07% and represented the variance between susceptible populations of *P. annua*. In InDels based PCA (Figure 7B), the PC1 and PC2 accounted for 18.36% and 11.40% variance, respectively. Variants were selected based on if they were present in all replicates of all indaziflam-resistant populations and absent in all susceptible populations. We found no InDel being of “HIGH” and/or “MODERATE” importance in this subset, while 192 SNPs satisfied these conditions. From the distribution of the SNPs across the genome, subgenome B had 1.6× more SNPs than A overall, in chromosomes 1, 2, 3, 4, and 7, with chromosome 7A having seven SNPs, whereas chromosome 7B had 18 SNPs (Figure 7C). We found several proteins involved in vesicle transport and intracellular trafficking that had SNPs causing amino acid changes. Some of the interesting genes identified with SNPs were coatomer γγ subunit, syntaxin 22 (a vesicle-associated membrane protein), and Ccz1 (a vacuolar fusion protein); all of which mediate intracellular trafficking in one way or another. The coatomer γγ subunit was especially interesting as it helps form the coatomer complex which in turn helps form vesicles that transport between the endoplasmic reticulum (ER) and Golgi (Duden, 2003). Syntaxin-22, a SNARE protein, is needed for vesicle merger from the ER to Golgi (Malsam and Söllner, 2011). Further, we found non-synonymous SNPs in proteins involved in cell wall modification, such as expansin-like A1, which causes cell-wall loosening, dirigent protein 9, for lignin synthase, as well as multiple glycosyl transferase proteins that are involved in cell carbohydrate metabolism and synthesis. Many transcription factors, including an RNA-binding superfamily, zinc finger proteins, and piwi, argonaute, and zwille (PAZ) domain proteins, had non-synonymous SNPs. Finally, some proteins involved in cytoskeleton and cell structure, such as actin-related protein, spiral1 protein, or formin-like proteins were also identified (Supplementary Table 9).

**Figure 7.**
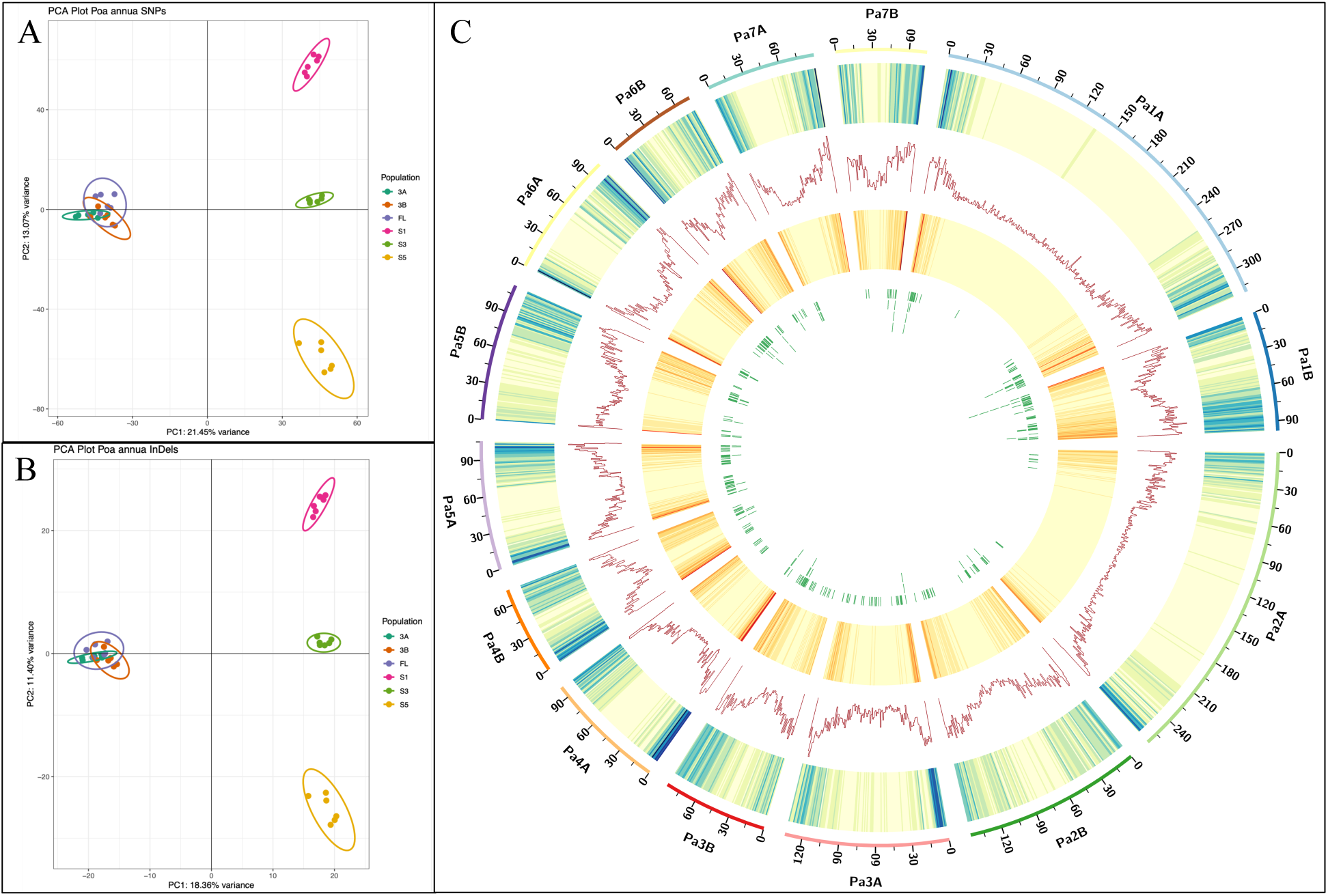
PCA plot of the variants across the resistant and susceptible populations, using (A) SNPs (B) InDels, suggesting potential single origin of the resistant biotypes, with distinct origins of the susceptible ones (C) Circos plot representation of variants and their distribution across the *P. annua* genome, From the outer to inner tracks: a) Chromosomes of *P. annua*, with A and B referring to A and B subgenomes (Mb), b) SNP density across the 1Mb genomic windows across all the populations, c) gene density per 1Mb genomic regions, d) indels density across the 1Mb genomic regions across all the populations e) High and moderate impact SNPs present only in resistant populations, represented as tiles across the genome.

Mahey et al. 2026 has reported vesicle trafficking as a putative target site of indaziflam with various coatomer subunits being recognized as top-candidates interacting with indaziflam (Mohit Mahey *et al*., in review). Further, the detection of SNP causing a Thr432Arg amino acid change in the coatomer subunit γγ, which has been consistently present in all the replicates of resistant populations but none in susceptible populations. Based on these initial results, we decided to take a closer look at the SNP causing a Thr432Arg amino acid change in the coatomer subunit γγ by exploring its presence in indaziflam-resistant *P. annua* populations identified recently in Oregon orchards (Miranda *et al*., 2026). Nanopore sequencing represented the average ratio of populations rather than individual plants, as ∼10 mg seed from each population were used in this analysis. The SNP causes a “C” in susceptible to mutate to “G” resulting in a threonine to arginine amino acid switch at the 432 position. This SNP was completely absent in the Oregon populations susceptible to indaziflam, while the G/C ratio varied among the resistant populations tested. This SNP was identified (in varied ratios) in several of the indaziflam-resistant populations from Oregon orchards (izr-4, izr-19, izr-5 and izr-18), although it was absent in a subset of resistant populations (izr-2 and izr-9) as well (Supplementary Figure 4).

## 4 DISCUSSION

*P. annua* is a notorious weed with evolved resistance to almost all available modes of action through both TSR and NTSR mechanisms. Recently, indaziflam resistance has become increasingly more prevalent in turfgrass systems and orchards, especially within *P. annua* populations resistant to other modes of action. WGCNA analysis of three susceptible and three indaziflam-resistant populations revealed a module that was positively correlated with elevated ED_50_ values. This module contained multiple P450s and GST metabolism genes. Of these, a CYP81A91_A was found to be overexpressed in all indaziflam- resistant populations of turfgrass systems (Table 2). Additionally, the CYP81A91_A gene had the highest MM and GS among all CYPs identified. Further, CYP81A91_A shared 92% sequence homology to a reported cytochrome P450 in *Lolium rigidum* that confers resistance to at least five herbicide chemistries (Han *et al*., 2021), including a triazine herbicide. The high sequence homology suggested that the overexpressed CYP81A91_A may be able to metabolize other small molecules containing triazine rings such as indaziflam. Cytochrome P450s are widely abundant, promiscuous enzymes in plants that have undergone rapid neo-functionalization and have emerged as the top gene families candidates involved in herbicide metabolism^78,79^. For example, cytochrome P450s identified in wheat (*Triticum aestivum;* CYP71) (Goldberg-Cavalleri *et al*., 2025) and rice (*Oryza sativa*; CYP81) (Ha *et al*., 2022) have been reported to metabolize several herbicides targeting ALS and PSII. Similarly, a cytochrome P450 from corn (*Zea mays*; CYP450) has been shown to metabolize herbicides from five modes of action including inhibitors of ALS, HPPD, PPO, and PSII, as well as a synthetic auxin (Brazier-Hicks *et al*., 2022).

Heterologous transformation of CYP81A91_A into *Arabidopsis thaliana* revealed no decreased sensitivity to indaziflam application on agar plates. However, these transgenic lines did show significant resistance to simazine, another triazine herbicide. Given what is known about P450 activity, we suspect this gene is removing one of the ethyl groups of the simazine, whereas for indaziflam, the triazane and indyl rings would likely need to fracture for resistance to manifest. In both cases, it is unclear whether these reactions alone would be enough to detoxify either herbicide. Often, detoxification requires a phase two reaction such as the conjugation of the herbicide to glutathione or a sugar (Gaines *et al*., 2020), albeit not always. Without the proper GST partner, our transgenic lines may not have had the required machinery to detoxify indaziflam, while other endogenous conjugating chemistries may be able to perform this operation for simazine in *A. thaliana*. Indeed, we did identify a GST in our highly correlated WGCNA module. This suggests the *A. thaliana* might lack the specific GST needed to perform indaziflam specific phase two metabolism. While this is one explanation for why we observed resistance to simazine and not indaziflam, it could also be that enzyme efficiency for indaziflam metabolism is low, requiring a level of expression we did not achieve in our heterologous system, or that expression in germinating roots does not protect the zone of cell division quickly enough upon germination.

Resistance to indaziflam may result from a combination of TSR and NTSR mechanisms. Whole transcriptome-wide SNP calling led to the identification of several non-synonymous mutations in multiple genes, some of which are involved in vesicle transportation and cell wall formation. These SNPs were unique in all indaziflam-resistant *P. annua* populations collected from golf course turfgrass in the southeastern United States and completely absent in all susceptible populations. Of these, the most interesting was a SNP causing an amino acid change of threonine to arginine at the 432 position (Thr432Arg) in the coatomer subunit γγ. This amino acid change turns the neutral threonine to a positively charged arginine. Coatomer proteins are involved in vesicle trafficking to and from the ER to the Golgi apparatus. Coatomer proteins play a crucial role in the recycling of the cellulose synthase enzyme, as well as the transfer of other proteins between organelles. The presence of this SNP could potentially cause less efficient binding of indaziflam, if it is indeed targeting the coatomer complex and not CESA directly. This, along with mild detoxification by the upregulated P450, may be what is ultimately conferring indaziflam- resistance in the *P. annua* populations included in this research. Interestingly, Mahey et al. 2026 reported the coatomer subunit γγ as one of the most enriched proteins in a pull-down assay conducted using indaziflam as bait (Mohit Mahey *et al*., in review).

To independently verify the involvement of Thr432Arg SNP in the coatomer subunit γγ in indaziflam-resistance, we screened indaziflam-resistant biotypes from orchards in Oregon for the presence of this SNP. The SNP was absent in all Oregon susceptible populations yet present in the majority of the indaziflam-resistant populations originating from orchards. Although the SNP was not present in all indaziflam-resistant populations tested from Oregon orchards, its absence in susceptible populations and detection in many of the most indaziflam-resistant populations analyzed is suggestive that it may be at least partially involved. The change in a single amino acid has been shown many times to cause herbicide resistance, as is the case for EPSPS-, ALS-, and ACCase-inhibiting herbicides (Whaley *et al*., 2007; Vila-Aiub *et al*., 2015; Han *et al*., 2017). We also found multiple other proteins with SNPs causing amino acid changes that are involved in cell wall and vesicle trafficking processes, each of which needs to be considered and tested in turn (Supplementary Table 9).

Understanding the resistance mechanism of *P. annua* is critical for identifying and tracking resistance and for making herbicide recommendations to practitioners. Indaziflam is an important herbicide for *P. annua* control and the speed at which resistance is evolving is alarming. Furthermore, evolution of NTSR by metabolism is especially concerning as these singular mechanisms can confer resistance to a large number of herbicides spanning modes of action and in unpredictable ways, meaning that even newly introduced herbicides may already be rendered ineffective. Our study showed that indaziflam-resistant *P. annua* has several cytochrome P450s upregulated, each of which probably contributes some level of resistance to a variety of herbicides. Heterologous transformation of the leading candidate (CYP81A91_A) conferred no clear resistance to indaziflam; however, it did provide significant resistance to another common *P. annua* herbicide, simazine. Further research needs to be done to untangle these putative overlapping resistance mechanisms, identifying which gene provides resistance to what set of herbicides, and the role that target site mutations such as Thr432Arg in coatomer subunit γγ play in these *P. annua* populations.

### SUPPORTING DATA

supplemental_poa_annua_indaziflam_v5.docs has

ST1: Quality metrics of raw RNA-seq data

ST2: Quality metrics of aligned reads

ST3: Hub genes of module firebrick3

ST4: Hub genes for module darkseagreen4

ST5: Hub genes of module green

ST6: ED_50_ values of the dry biomass of *P. annua* populations

ST7: Alignment statistics of pairwise alignment of *Poa annua* CYP81A91_A with *Lolium rigidum* CYP81A10v7

ST8: Delta-delta Cq analysis of transformed CYP81A91_A

ST9: candidate SNPs identified

SF1: Copy number assay of the transformed CYP81A91_A in *A. thaliana*

SF2: phenotype of indaziflam dose response of three independent transformed *A. thaliana*

SF3: The phenotype of the transformed plants with simazine

SF4: Coatomer gamma SNP ratio among various Oregon populations

## ACKNOWLEDGEMENTS

The authors would like to thank Luan Cutti, Jan Michals, Nash Hart, and Ednaldo Borgato for their valuable insights in the greenhouse study. Thank you to Bayer Environmental Sciences for funding to support the RNA sequencing.

## AUTHOR CONTRIBUTIONCONTRIBUTIONS

MM, PKL, ELP: planned and designed research, MM, MO: performed experiments, greenhouse work, MM, MO, JTB, PKL, ELP: analyzed data, MM: Bioinformatics analysis, MM: writing – original draft, MM, MO, JTB, PKL, ELP: writing – review and editing, JMT, MLM, JTB: provided genotype, ELP: funding acquisition, PKL,ELP: resources

## DATA AVAILABILITY STATEMENT

The RNA-seq data are available at through NCBI at project number PRJNA1465323. All the scripts used for analysis are available at GitHub: https://github.com/mohitmahey/RNA_seq_analysis and https://github.com/mohitmahey/whole_transcriptome_variant_calling

